# Phylogenetically based establishment of a dengue virus panel, representing all available genotypes, as a tool in dengue drug discovery

**DOI:** 10.1101/439695

**Authors:** Franck Touret, Cécile Baronti, Olivia Goethals, Marnix Van Loock, Xavier de Lamballerie, Gilles Querat

## Abstract

Dengue fever is the most widespread of the human arbovirus diseases, with approximately one third of the world’s population at risk of infection. Dengue viruses are members of the genus *Flavivirus* (family *Flaviviridae*) and, antigenically, they separate as four closely related serotypes (1-4) that share 60 to 75 % amino acid homology. This genetic diversity complicates the process of antiviral drug discovery. Thus, currently no approved dengue-specific therapeutic treatments are available. With the aim of providing an efficient tool for dengue virus drug discovery, a collection of nineteen dengue viruses, representing the genotypic diversity within the four serotypes, was developed. After phylogenetic analysis of the full-length genomes, we selected relevant strains from the EVAg collection at Aix-Marseille University and completed the virus collection, using a reverse genetic system based on the infectious sub-genomic amplicons technique. Finally, we evaluated this dengue virus collection against three published dengue inhibitory compounds. NITD008, which targets the highly conserved active site of the viral NS5 polymerase enzyme, exhibited similar antiviral potencies against each of the different dengue genotypes in the panel. Compounds targeting less conserved protein subdomains, such as the capsid inhibitor ST-148, or SDM25N, a ∂ opioid receptor antagonist which indirectly targets NS4B, exhibited larger differences in potency against the various genotypes of dengue viruses. These results illustrate the importance of a phylogenetically based dengue virus reference panel for dengue antiviral research. The collection developed in this study, which includes such representative dengue viruses, has been made available to the scientific community through the European Virus Archive to evaluate novel DENV antiviral candidates.

## Introduction

Dengue virus (DENV) is a major threat to human health, with approximately one third of the world’s population at risk of being infected. DENV is the causative agent of dengue fever, as well as the more severe dengue haemorrhagic fever (DHF)^1^ and dengue shock syndrome (DSS). It belongs to the genus *Flavivirus* (*Flaviviridae* family), which comprises other clinically important human pathogens, such as yellow fever virus, West Nile virus and the recently emerging Zika virus^2^. DENV is an arthropod borne virus transmitted through the bite of infected mosquitoes from the genus *Aedes* (Stegomyia). Epidemiological transmission of DENV is confined to urban and peri-urban cycles for which *Aedes aegypti* and *Ae albopictus* mosquitoes, respectively, are the primary transmission vectors^3^. Dengue is a positive-sense single stranded RNA virus with a 10.7 kb genome encoding a single polyprotein which is post-translationally processed into three structural proteins, viz., capsid (C), pre-membrane (prM), envelope (E) and seven non-structural proteins (NS1, NS2A, NS2B, NS3, NS4A, NS4B and NS5)^4^. Four antigenically closely related serotypes of DENV (1-4) which share 60 to 75 % amino acid homology, have been identified^5^. Within this serotype demarcation, the DENV are also grouped into genotypes, with varying terminology between authors^6,7^ (hereunder we refer to the grouping proposed by Weaver and Vasilakis^7^).

Hence, many of the DENV diagnostic tools do not readily distinguish between DENV serotypes. Moreover, co-circulation of different serotypes during DENV epidemics^8^ increases the complexity of virus identification. Added to these factors, antibody dependent enhancement of the disease i.e., when patients contract a heterotypic secondary DENV infection^9^ is a potential additional complication for effective treatment of patients. Consequently, scientists are faced with the challenge of developing Directly Active Antivirals (DAA) that can inhibit the entire spectrum of genetically diverse serotypes and/or genotypes of DENV. However, despite the tireless efforts to provide an antiviral therapy^10–13^, there are still no approved drugs on the market to treat dengue infections. At present, the treatments available are merely supportive^14^.

A major barrier to evaluating the activity spectrum of potential DENV-inhibitory molecules arises from the non-availability of a well-defined panel of viruses that specifically represents the genetic variability of all characterised DENV isolates. With the aim of providing a tool for DENV research, with which to assess the antiviral activity of potential inhibitory molecules, we have developed a collection of DENV with sequences that include representative genotypes from within the four DENV serotypes (figure 1). Wherever possible, we selected clinical strains with a limited number of passages in cell culture. Strains were selected from either the European Virus Archive (EVA) collection^15^, the French National Reference Centre for arboviruses (CNR), or the World Reference Center for Emerging Viruses and Arboviruses (WRCEVA). Viruses that could not be obtained but for which full length genome sequences were available, were re-created using the versatile infectious sub-genomic amplicons (ISA) reverse genetics technology^16,17^.

**Figure 1:**
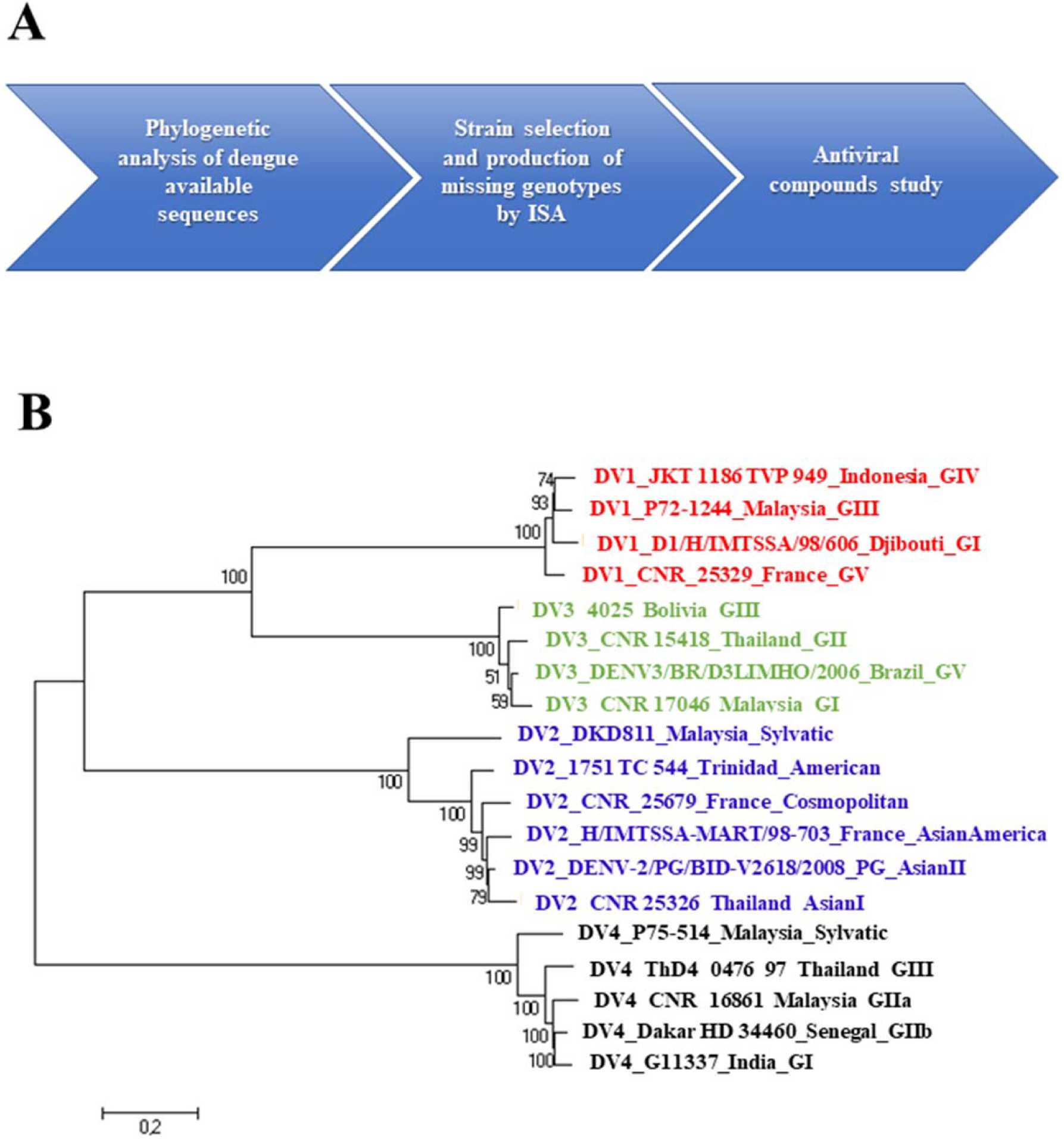
Global serotype-representative DENV collection. A: Pipeline of the workflow employed for the virus collection. B: Maximum likelihood phylogenetic tree (GTR+G+I model with 500 bootstraps), based on the complete nucleic acid sequences of the virus collection. Strain information: DENV-1: Genotype I: (Djibouti, 1998, AF298808); Genotype III (Malaysia,1972, EF457905.1); Genotype IV (Indonesia, 1977, EUO74031); Genotype V (France, 2014, MF004384); DENV-2: Genotype Asian-America (France Martinique, 1998, AF208496); Genotype American (Trinidad, 1953, EU073981.1); Genotype Cosmopolitan (France, 2014, MF004385); Genotype Asian 1 (Thailand, 2014, MH888331); Genotype Asian 2 (Papua New Guinea, 2008, FJ906959.1); Genotype Sylvatic (Malaysia, 2008, FJ467493.1); DENV-3: Genotype I (Malaysia, 2012, MF004386); Genotype II (Thailand, 2012 MH888332); Genotype III (Bolivia,2011, MH888333); Genotype V (Brazil,2006, JN697379.1); DENV-4: Genotype IIb (Senegal, 1981, MF004387,); Genotype IIa (Malaysia, 2013, MH888334); Genotype III (Thailand, 1997, AY618988.1); Genotype Sylvatic (Malaysia, 1975, JF262779.1); Genotype I (INDIA, 1961, JF262783.1). Complete information relevant to the strains of the collection are more fully detailed in the supplemental material.

In order to select representative genotypes, we collected dengue full-length genome sequences from the NCBI database and complemented this database with those of our, still unpublished, “in house” and CNR strains. We performed phylogenetic reconstructions with the maximum likelihood method to assign all available genome sequences to a genotype in a serotype (supplementary material Fig 1, 2, 3 and 4). Within each genotype, we focused on strains that were not subjected to extensive cell passage and were either available as biological isolates in virus collections or as full-length sequences in GenBank. Six dengue genotypes were available only as complete genome sequences in the NCBI database without any biological strain counterparts in referenced collections (DENV-1 genotype III, DENV-2 genotype sylvatic and Asian II, DENV-3 genotype V and DENV-4 genotype III and sylvatic). Two genotypes were not available at all because of incomplete genome sequence (DENV-1 genotype II and DENV-3 genotype IV). To obtain the biological viruses from the completely sequenced strains, we designed reverse genetics systems based on the ISA technique^16–18^ and generated synthetic overlapping DNA fragments that covered each of the entire genome, bordered by a CMV promoter on the 5’ end and a Ribozyme and poly-adenylation signal on the 3’ end. The overlapping fragments were co-transfected into a mix of human and hamster embryonic kidney cell lines (HEK 293 and BHK-21 purchased from the American Cell Culture Collection). This enabled us to recover the missing biological strains to complete the collection. The initial viral stocks were amplified in Vero E6 cells and fully sequenced. All the DENV strains used, have been made available through the EVAg collection (https://www.european-virus-archive.com/).

Various specific dengue inhibitors that target several viral proteins involved in different replication steps, have been discovered. ST-148, an inhibitor targeting the capsid structural protein, has been reported to inhibit all DENV serotypes in cell culture, although with varying efficiency. This inhibitor also appears promising in the AG-129 mouse model when infected with a strain of DENV-2^19^. NITD008, an adenosine analogue inhibitor that targets the RNA-dependent RNA polymerase activity, was shown to be inhibitory against all dengue serotypes as well as other flaviviruses, including West Nile virus, yellow fever virus and tick-borne Powassan virus^13^. SDM25N, a ∂ opioid receptor antagonist, has been reported to target the NS4B protein, probably indirectly through a cellular factor. Thus far, it has only been shown to be active against a DENV-2 strain^12^. Based on the different mechanisms of action of the 3 compounds, their respective target and its associated sequence variability across the different genotypes, we hypothesize that the antiviral activity of the compounds might differ between all of the genotypes of DENV. Therefore, the antiviral activity of these three compounds was assessed using a single common protocol based on a viral RNA yield reduction assay^20^. The assay did not depend on the cytopathogenic potential of the strain, thus allowing for the inclusion of any dengue strain in the panel tested. The compounds were assayed from 10 to 0.004µM, with 3-fold step-dilution in triplicate. The amount of viral RNA in the supernatant medium, sampled at intervals during the growth cycle, was quantified by qRT-PCR to determine the 50% maximal effective concentration (EC_50_) (Table 1).

**Table 1.** Dengue virus collection-susceptibility to three antiviral compounds assessed by yield reduction assay. Anti-capsid ST-148, nucleoside analogue NITD008 and ∂ opioid receptor antagonist SDM25N were independently tested twice, with 3 replicates per experiment, against the dengue collection from 10µM to 0.005µM. AA: Asian American, A: American, C: cosmopolitan.

|  |  |  | ST-148 | NITD008 | SDM25N |
| --- | --- | --- | --- | --- | --- |
|  | Virus | Genotype | EC50 (µM) | EC50 (µM) | EC50 (µM) |
| **Dengue 1** | D1/H/IMTSSA/98/606<br>**Djibouti** | **I** | **>10** | **0,9±0,1** | **>10** |
|  | JKT 1186 TVP 949<br>**Indonesia** | **IV** | **>10** | **0,3±0,03** | **5,5±3,67** |
|  | CNR_25329<br>**France** | **V** | **>10** | **2,7±4** | **7,4±0,04** |
|  | P72-1244<br>**Malaysia** | **III** | **3±0,5** | **0,9±0,2** | **>10** |
| **Dengue 2** | H/IMTSSA-MART/98-703<br>**France** | **AA** | **0,8 ±0,5** | **0,9±0,3** | **2,9±0,95** |
|  | _1751 TC 544<br>**Trinidad** | **A** | **1 ±0,7** | **0,3±0,06** | **2,9±0,01** |
|  | CNR_25679<br>**France** | **C** | **1,1 ±0,3** | **0,2±0,07** | **1,9±0,03** |
|  | CNR 25326<br>**Thailand** | **Asian I** | **0,1 ±0,03** | **0,9±0,2** | **7,7±0,04** |
|  | DENV-2/PG/BID-V2618/2008<br>**Papua New Guinea** | **Asian II** | **0,2±0,16** | **0,3±0,5** | **4,1±0,02** |
|  | DKD811<br>**Malaysia** | **Sylvatic** | **0,4 ±0,18** | **0,4±0,1** | **>10** |
| **Dengue 3** | DENV3/BR/D3LIMHO/2006<br>**Brazil** | **V** | **>10** | **1±0,09** | **>10** |
|  | 4025<br>**Bolivia** | **III** | **>10** | **1±0,05** | **>10** |
|  | CNR 17046<br>**Malaysia** | **I** | **>10** | **2,8±0,3** | **>10** |
|  | CNR 15418<br>**Thailand** | **II** | **>10** | **1,2±0,3** | **>10** |
| **Dengue 4** | G11337<br>**India** | **I** | **>10** | **1,2±0,03** | **>10** |
|  | Dakar HD 34460<br>**Senegal** | **IIb** | **>10** | **0,9±0,3** | **>10** |
|  | CNR_16861<br>**Malaysia** | **IIa** | **>10** | **0,4±0,01** | **>10** |
|  | ThD4_0476_97<br>**Thailand** | **III** | **0,3±0,08** | **0,2±0,08** | **>10** |
|  | P75-514<br>**Malaysia** | **Sylvatic** | **>10** | **1±0,05** | **>10** |

The DENV strains of the collection showed similar sensitivity towards the nucleoside analogue inhibitor NITD008 with EC_50_’s ranging from 0.2µM to 2.8µM, which is in accordance with previously published results^21^.

The capsid inhibitor ST-148 inhibited all DENV-2 genotypes with EC_50_’s ranging from 0.25 to 1.1µM. However, only one genotype for DENV-1 (DENV-1 GIII at 0.5µM), and for DENV-4 (DENV-4 GIII at 0.3µM), were inhibited by this compound. Finally, no activity was observed against DENV-3 genotypes, with EC_50_’s > 10µM. Although Byrd and co-workers^19^ found that the DENV-2 serotype was the most sensitive serotype to this capsid inhibitor and showed some variability in the inhibition against other serotypes, they did not fully evaluate the variation in susceptibility to other serotypes sufficiently comprehensively to draw conclusions.

SDM25N showed moderate efficacy, EC_50_’s ranging from 1.7 – 7.7 µM against a large proportion of the DENV-2 genotype strains, and half of the DENV-1 genotypes. However, no activity was observed against none of the DENV-3 and 4 genotypes with EC_50_’s above 10 µM. This result suggests that the binding affinity of NS4B to the hypothetical cellular factor targeted by SDM25N varies greatly among various DENV genotypes and/or that this cellular factor might be dispensable for efficient replication of some DENV genotypes.

Overall, the results demonstrate that compounds targeting highly conserved sites, exemplified by nucleoside analogue inhibitor NITD008 (targeting the active site of the polymerase), had a broader pan-serotypic activity, with similar EC_50_’s regardless of the DENV genotype. In contrast, compounds targeting less conserved proteins or protein subdomains, either directly (*e.g.* the capsid) or indirectly through an interaction with a host factor of the cell (*e.g.* SDM25N), exhibited larger differences in activity towards the various genotypes of DENV.

Importantly, these data illustrate the fact that a sound *in cellulo* evaluation of anti-dengue candidate molecules requires the use of a complete reference virus panel that enables estimates of the antiviral activity against each of the identified DENV genotypes to be obtained. Modern reverse genetics techniques have enabled us to develop such a representative collection, and it has been made available to the scientific community through the European Virus Archive collection (EVA). We believe that the availability of this new tool will enable the independent assessment of pan-serotypic activity of anti-dengue candidates in the future, fulfilling a critical requirement for a successful dengue antiviral small molecule.

## Acknowledgments

We would thank Dr Robert Tesh from UTMB (Texas) for providing the dengue 4 Genotype I strain G11337 from India. We thank Dr Raphaelle Klitting for her help and expertise regarding the dengue phylogeny. We would thank Pr E.A Gould and Dr Stéphane Priet for their careful reading of the manuscript. As well as Magali Gilles, and Fiona Baudino from the UMR UVE (Marseille, France) for excellent technical help. We also thank Geraldine Piorkowski and Karine Barthelemy from UMR UVE (Marseille, France) for the sequencing. This research was carried out under sponsorship of Janssen-Cilag S. A., a pharmaceutical company of Johnson & Johnson (Research agreement ICD#1041950) and EVAg European Union – Horizon 2020 programme under grant agreement no. 653316; http://www.european-virusarchive.com).

## Contributions

OG, MVL, GQ and XDL generated the idea of the panel. FT, XDL and GQ conceived the experiments. XDL proposed the study design. FT, CB, and GQ performed the experiments. FT and GQ analyzed the results. FT and GQ wrote the paper. FT, CB, GQ, OG, MVL and XDL reviewed and edited the paper.

