## Supplemental data for "Phylogenetically based establishment of a dengue virus panel, representing all available genotypes, as a tool in dengue drug discovery"

[1] : Unité des Virus Émergents (UVE : Aix-Marseille Univ – IRD 190 – Inserm 1207 – IHU Méditerranée Infection), Marseille, France

[2] : Janssen Infectious Diseases Discovery, Janssen Pharmaceutica NV, Turnhoutseweg 30, 2340, Beerse, Belgium.

**Supplemental Table 1** Comparison of the antiviral susceptibility between a clinical strain and a strain obtained by ISA**.**

| | Dengue 1<br>D1/H/IMTSSA/98/606<br>**Djibouti** | | Dengue 3<br>CNR 17046<br>**Malaysia** | | Dengue 4<br>Dakar HD 34460<br>**Senegal** | |
|---|---|---|---|---|---|---|
| **Origin** | **Clinic** | **ISA** | **Clinic** | **ISA** | **Clinic** | **ISA** |
| **NITD008 EC50 (µM)** | **0,9** | **0,54** | **2,8** | **3,57** | **0,9** | **0,55** |
| **ST-148 EC50 (µM)** | **>20** | **>20** | **10,66** | **19,17** | **12,87** | **17,66** |
| **SDM25N EC50 (µM)** | **7,28** | **17,3** | **18,8** | **>20** | **8,8** | **11,9** |

### Supplemental Material and methods

#### Cell lines

All cells were grown at 37 °C with 5 % CO 2 with 1 % penicillin/streptomycin (PS; 5000 U ml −1 and 5000 µg ml −1; Life Technologies) and supplemented with 1% non-essential amino acids (Life Technologies) in specific cell line medium.

Baby Hamster kidney (BHK21 ATCC) and Human Embryonic kidney cells (HEK293 ATCC) cells were grown in Dulbecco's Modified Eagle's Medium High glucose (4500 mg/l) (Life Technologies) with 7.5 % heat-inactivated fetal bovine serum (FBS; Life Technologies).

C636 cells (ATCC) were grown at 28 °C Leibovitz 15 Media (Life Technologies)

VeroE6 (ATCC) cells minimal essential medium (Life Technologies) with 7.5 % heat-inactivated fetal bovine serum (FBS; Life Technologies).

**Viruses**

**DENV-1 strains**: Genotype I: (Djibouti, strain D1/H/IMTSSA/98/606, GenBank: AF298808, EVAg *Ref-SKU*: 001V-02338); Genotype II: not available, no complete genome sequence; Genotype III (Malaysia, strain P72-1244, GenBank EF457905.1, EVAg *Ref-SKU*: *001V-03105*); Genotype IV (Indonesia, strain JKT 1186 TVP 949, GenBank: EUO7031, EVAg *Ref-SKU*: 001V-02337); Genotype V (France, CNR_25329, GenBank MF004384, EVAg *Ref-SKU*: 001V-02228);

**DENV-2 strains**: Genotype Asian/America (Martinique, strain H/IMTSSA-MART/98-703, GenBank: AF208496, EVAg *Ref-SKU*: 001v-EVA1019); Genotype American (Trinidad, strain Trinidad 1751, GenBank: EU073981.1, EVAg *Ref-SKU*:001v-EVA928; Genotype Cosmopolitan (France, CNR_25679, GenBank MF004385, EVAg *Ref-SKU*: 001V-02229); Genotype Asian I (Thailand, CNR_25326, GenBank MH888331 EVAg *Ref-SKU*: *001V-02340*); Genotype Asian II (Papua New Guinea, strain BID-V2618, GenBank FJ906959.1, EVAg *Ref-SKU*: *001V-03106*); Genotype Sylvatic (Malaysia, strain DKD811, GenBank FJ467493.1, EVAg *Ref-SKU*: *001V-03107*);

**DENV-3 strains**: Genotype I (Malaysia, CNR_17046, GenBank MF004386, EVAg *Ref-SKU*: *001V-02230*); Genotype II (Thailand, strain CNR_15418, GenBank MH888332EVAg *Ref-SKU*: *001V-02296*); Genotype III (Bolivia, strain BOL_4025, GenBank MH888333 EVAg *Ref-SKU*: *001v-EVA1477*); Genotype IV: not available, no complete genome sequence; Genotype V (Brazil, strain DENV3/BR/D3LIMHO/2006, GenBank JN697379.1, EVAg *Ref-SKU*: *001V-03108*);

**DENV-4 strains**; Genotype I (strain G11337, India, GenBank JF262783.1, kindly provided by UTMB); Genotype IIa (Malaysia, strain CNR_16861, GenBank MH888334, EVAg *Ref-SKU*: *001V-02290*); Genotype IIb (Senegal, strain Dak HD 34 460, GenBank MF004387, EVAg *Ref-SKU*: *001v-EVA930*); Genotype III (Thailand, strain ThD4_0476_97, GenBank AY618988.1 EVAg *Ref-SKU: 001V-03109*); Genotype Sylvatic (Malaysia, strain P75-514, GenBank JF262779.1, EVAg *Ref-SKU: 001V-03110*);

To prepare our working stock a 25 cm2 culture flask of confluent Vero E6 cells growing with MEM medium with 2.5 % FBS (Life Technologies) was inoculated with 100 µl of infectious supernatant. Cell supernatant medium was harvested at the peak of infection and supplemented with 25mM HEPES (Sigma) before being stored freeze in small aliquots at -80°C.

All experiments with replicative viruses were performed in BSL3 laboratory

All virus strains are available for the community at European Virus Archive. https://www.european-virus-archive.com/

**DNA preparation**

PCR amplification of *de novo* synthetized gene (Genscript) was done using Platinum PCR SuperMix High Fidelity kit (Life Technologies), in a final volume of 50 µl (primers used for reverse genetic are listed in table 2). Amplifications were performed on a thermal cycler ABI 9700 (Applied biosytem) with the following conditions: 94 °C for 2 min followed by 40 cycles of 94 °C for 15 s, 60 °C for 30 s, 68 °C for 5 min plus 10 min final elongation at 68 °C. Then the CMV promoter (pCMV) and the hepatitis delta ribozyme followed by the simian virus 40 poly-adenylation signal (HDR/SV40pA) were added by fusion PCR at, respectively the 5' and 3' end of the first and the third DNA fragment. Briefly the pCMV was amplified by PCR from a previously available plasmid [1] using a forward specific primer and a reverse primer extended with the first seventy nucleotides of the corresponding dengue viral genome. This tagged pCMV and the first DNA fragments were then merged by fusion PCR. The HDR/SV40pA was amplified by PCR from a previously available plasmid [1] using a forward primer extended with the last 70 nucleotides of the corresponding dengue viral genome. This tagged HDR/SV40pA and the third DNA fragment were then merged by fusion PCR. Fusion PCR was performed using same master mix and condition as described above. PCR fragments size and quality were verified by running gel electrophoresis and PCR products were directly purified using the High pure PCR product purification kit (Roche) avoiding an agarose gel purification step.

**Supplemental table 2:** Primers used in this study to generate different dengue by reverse genetic using the ISA technic.

| Strains | Fragment | Primer | Sequence |
|---|---|---|---|
| DENV-1 Genotype 3 EF457905.1 | F1 | DVA1F | AGTTGTTAGTCTGTGTGGACCGAC |
| | | DV1G3 A1R | TGAGAGGTCGTAGACAGGCA |
| | F2 | DV1G3 A2F | CAGTCACATCAGTTATGGGC |
| | | DV1G3 A2R | CTAGGCCTCTTCCTGATAGG |
| | F3 | DV1G3 A3F | CCTACAGACACGCTATGGAAGAG |
| | | DVA3R | AGAACCTGTTGATTCAACAGCACC |
| | CMV-F1 | DV1G3A1Rint | AGGTCGTAGACAGGCATAGAG |
| | | DV1G3-CMV70R | ACTGTTAGAACTACGTTAAGCAAGCTTCCGATTCGAAACTGTTCTTGTCGGTCCACGTAGACTAACAACTCGGTTCACTAAACGAGCTCTGCTT |
| | F3-HDR/SV40pA 70 | DV1G3A3fint | AGACATGCAATGGAAGAACTCC |
| | | DV1G3-SV70F | TGTCTCTACAGCATCATTCCAGGCACAGAACGCCAGAAAATGGAATGGTGCTGTTGAATCAACAGGTTCTGGCCGGCATGGTCCCAGC |
| DENV-2 Genotype Asian2 FJ906959 | F1 | DVA1F | AGTTGTTAGTCTGTGTGGACCGAC |
| | | DV2A2 A1R | CTCTGGTATGGTGCTCTGGG |

| | | | |
|---|---|---|---|
| | F2 | DV2A2 A2F | GCAGCCTTCAAAGTCAGACC |
| | | DV2A2 A2R | AAAGATTCCTCCTGTGACTGTAGC |
| | F3 | DV2A2 A3F | AACTTAGCAGTGCTGCACAC |
| | | DVA3R | AGAACCTGTTGATTCAACAGCACC |
| | CMV-F1 | DV2A2 A1Rint | TATGGTGCTCTGGGAGAGG |
| | | DV2A2-CMV70R | CTGTTAGAACTACGTTGAGCTTAGCTCCCTCAAAGAATCTGTCTTTGTCGGTCCACGTAGACTAACAACTCGGTTCACTAAACGAGCTCTGCTT |
| | F3-HDR/SV40pA 70 | DV2A2 A3Fint | GTGCTGCACACGGCTGA |
| | | DV2A2-SV70F | TGTCTCCTCAGCATCATTCCAGGCACAGAACGCCAGAAAATGGAATGGTGCTGTTGAATCAACAGGTTCTGGCCGGCATGGTCCCAGC |
| DENV-2 Genotype Sylvatic FJ467493 | F1 | DVA1F | AGTTGTTAGTCTGTGTGGACCGAC |
| | | DV2GS A1R | AACTGTTTCTGGCATGTTGCT |
| | F2 | DV2GS A2F | ACCTTCAAAGTTAGGCCCACC |
| | | DV2GS A2R | CTAGTTGTTGTGTTGTGCCACC |
| | F3 | DV2GS A3F | TCTCATAGTGCTGCTCATCCC |
| | | DVA3R | AGAACCTGTTGATTCAACAGCACC |
| | CMV-F1 | DV2GS A1Rint | TAGTGCTGCTCATCCCTGAG |
| | | DV2GS-CMV70R | TGTTAGAACTACGTTAAGCCAAGCTTCCTCAAAGAAACTGTTCTTTGTCGGTCCACGTAGACTAACAACTCGGTTCACTAAACGAGCTCTGCTT |
| | F3-HDR/SV40pA 70 | DV2GS A3Fint | TAGTGCTGCTCATCCCTGAG |
| | | DV2GS-SV70F | TGTCTCCTCAGCATCATTCCAGGCACAGAACGCCAGAAAATGGAATGGTGCTGTTGAATCAACAGGTTCTGGCCGGCATGGTCCCAGC |
| DENV-3 Genotype 5 JN697379.1 | F1 | DVA1F | AGTTGTTAGTCTGTGTGGACCGAC |
| | | DV3G5 A1R | CTCTAGATGTCAGTTTCCTCAGGA |
| | F2 | DV3G5 A2F | CTCTCAGGGCAAATAACATGG |
| | | DV3G5 A2R | TTCCATCGTTTCTGGTAGTTCCT |
| | F3 | DV3G5 A3F | TGCCTTCACACTTAGCCCAC |

| | | | |
|---|---|---|---|
| | | DVA3R | AGAACCTGTTGATTCAACAGCACC |
| | CMV-F1 | DV3G5 A1Rint | AGATGTCAGTTTCCTCAGGAAGAAT |
| | | DV3G5-CMV70R | ACTGTCAGCACTACGTTAAGCAAGCTTCCGAGTCGAAACTGTTCTTGTCGGTCCACGTAGACTAACAACTCGGTTCACTAAACGAGCTCTGCTT |
| | F3-HDR/SV40pA 70 | DV3G5 A3Fint | ACACTTAGCCCACAGAACGA |
| | | DV3G5-SV70F | TGTCTCCTCAGCATCATTCCAGGCACAGAACGCCAGAAAATGGAATGGTGCTGTTGAATCAACAGGTTCTGGCCGGCATGGTCCCAGC |
| DENV-4 Genotype 3 AY618988.1 | F1 | DV4A1F | AGTTGTTAGTCTGTGTGGACCGAC |
| | | DV4G3 A1R | TGTTTCTCTTGAGGTGAGTTTCCT |
| | F2 | DV4G3 A2F | CGAGCCATTATCATGCTAGG |
| | | DV4G3 A2R | ATGAGTGTTTCCAATGACTCC |
| | F3 | DV4G3 A3F | CCTTGACAATATAGTCATGCTCCA |
| | | DV4A3R | AGAACCTGTTGGATCAACAAC |
| | CMV-F1 | DV4G3 CMV70R | ACTGTTAGAACTGTGTTAAGCAAGCTTCCGATTTGGAACTGTCCTCGTCGGTCCACACAGACTAACAACTCGGTTCACTAAACGAGCTCTGCTT |
| | | DV4G3 A1Rint | TCTTGAGGTGAGTTTCCTTAGGAA |
| | F3-HDR/SV40pA 70 | DV4G3 SV70F | GGCCGGCATGGTCCCAGCTGTCTCTGCAACATCAATCCAGGCACAGAGCGCCGCAAGATGGATTGGTGTTGTTGATCCAACAGGTTCT |
| | | DV4G3 A3Fnt | CAATATAGTCATGCTCCACACAAC |
| DENV-4 Genotype Sylvatic JF262779.1 | F1 | DV4A1F | AGTTGTTAGTCTGTGTGGACCGAC |
| | | DV4GS A1R | GTCATTGCCATTCCTATCACCA |
| | F2 | DV4GS A2F | CATGTTGGGAGATACCATGTTTGG |
| | | DV4GS A2R | TGTCTCCAATGATTCTGGGAGC |
| | F3 | DV4GS A3F | CTTGACAACATAGTCATGCTACAC |
| | | DV4A3R | AGAACCTGTTGGATCAACAAC |
| | CMV-F1 | DV4GS CMV70R | ACTGTTAGAACTGTGTTAAGCAAGCTTCCGATTCGGAACTGTCCTTGTCGGTCCACACAGACTAACAACTCGGTTCACTAAACGAGCTCTGCTT |

| | | | |
|---|---|---|---|
| | | DV4GS A1Rint | ATTGCCATTCCTATCACCATCAG |
| | F3-HDR/SV40pA 70 | DV4GS SV70F | GGCCGGCATGGTCCCAGCTGTCTCTGCAAC ATCAATCCAGGCACAGAGCGCCGCAAGATG GATTGGTGTTGTTGATCCAACAGGTTCT |
| | | DV4GS A3Fint | CAACATAGTCATGCTACACACAAC |
| pcMV | CMV-F1 | pCMV F | GAATAAGGGCGACACGGAAATGT |
| HDR/SV40pA | F3-HDR/SV40pA 70 | HDR/SV40pA R | AGCGGGTGTTTTTCCGAGTC |

**Virus production by ISA**

Virus were produced by reverse genetic as previously described [1–3]. Briefly, one-day prior transfection, a mix of BHK21 and HEK293 cells were seeded on 6 wells or 96 wells amine pure coat plate (Corning) to be sub confluent the next day. Two µg or 200 ng of an equimolar mix of the three fragments (sub genomic amplicons from the DNA preparation) were transfected using lipofectamine 3000 (Life Technologies), following manufacturer instruction, in 5 or 15 replicates. 6 days later, the supernatant was passed on a regular 6 or 96 well plate (Greiner) containing C6/36 cells. Then 4 to 6 days after infection viral supernatant was collected analyzed by qRT-PCR to assess viral replication.

**RNA extraction and virus quantification**

100µl of viral supernatant is collected into a S-Block (Qiagen) previously loaded with VXL lysis buffer containing proteinase K and RNA carrier. RNA extraction was performed using the Qiacube HT automat and the Cador Pathogen 96 HT kit to which we have added a DNAse (QIAGEN) digestion step, following manufacturer instruction. Viral RNA was quantified by real-time RT qPCR (EXPRESS One-Step Superscript™ qRT-PCR Kit, universal Invitrogen using 3.5 µL of RNA and 6.5 µL of RT qPCR mix and standard fast cycling parameters, i.e., 10 min at 50 °C, 2 min at 95 °C, and 40 amplification cycles (95 °C for 3 sec followed by 30sec at 60 °C). The four control wells of four 2 log dilutions of an appropriate T7-generated RNA standard of known quantities (100 copies to $100\times10^6$ copies) were include. RT qPCR reactions were performed on QuantStudio 12K Flex Real-Time PCR System (Applied Biosystems) and analyzed using QuantStudio 12K Flex Applied Biosystems software v1.2.3 The primers and probes used for the different serotypes during RT-qPCR are described in Table 3).

**Genome sequencing**

In order to verify our genome integrity after ISA and in our stock [3]. We then performed Reverse transcriptase PCR (Superscript III One-Step RT-PCR Platinum TaqHifi kit; Life Technologies) using the same set of primers that we used to generate the sub-genomic amplicons, except for the 5' and 3' extremities (primers used for sequencing are listed in table 1 and annotated seq). Then the complete resulting genome was pooled and analyzed using the Ion PGM Sequencer (Life technologies). Resulting reads were then analyzed using the CLC

Genomics Workbench 8 software. They were trimmed using quality score, by removing the previously described primers by systematically removing 6 nt at the 5′ and 3′ termini and finally by discarding those with under 30 nt length. Remaining reads were mapped using the designed sequence

#### Compounds Tested

SDM25N compound NS4B inhibitor [4], NITD008 nucleoside inhibitor [5]; Anti-capsid ST148 [6] were obtained from Hit2lead (www.hit2lead.com).

#### Antiviral Assay

One day prior to infection, $5\times10^4$ Vero E6 cells were seeded in 100 µl assay medium (containing 2.5 % fetal calf serum [FCS]) in 96-well plates. The next day, eight 3 fold serial dilutions of NITD008, ST-148 and SDM25N (0.004 µM – 10 µM), in triplicates were added to the cells (25 µl/well, in 2.5 % FCS-containing medium). Four virus control wells (per virus) were supplemented with 25 µL medium and four cell control wells were supplemented with 50 µl of medium. After 15 min, 25 µl of a virus mix diluted in medium was added to the wells at the correct MOI, determined so that the replication growth is still in the log growth curve for the readout at day 3 or 4, depending on the virus strain. Plates were incubated for 3 to 5 days at 37°C after which 100 µl of the supernatant was collected for viral RNA purification as described above in the RNA extraction and virus quantification paragraph.

**Supplemental table 3:** Primers and probes used for RT-qPCR

| Primer/probe | Sequence (5' - 3') | Target |
|---|---|---|
| DV-1 F5 | CRAGATGTCCRACACAAGGA | DENV-1 |
| DV-1 R5 | CGKCGACACACACAAARTTCG | DENV-1 |
| DV-1 P5 | ***FAM***-CTGGTGGAAGAACAAG-***MGB*** | DENV-1 |
| DV-2 F2 | TGGCAGCAATCCTGGCATA | DENV-2 |
| DV-2 R2 | GTCATTGAAGGAGCGACAGCT | DENV-2 |
| DV-2 P2 | ***FAM***-CRATAGGAACGACACATT-***MGB*** | DENV-2 |
| DV-2 A2P | ***FAM***-CCATAGGAACGACACATT ***MGB*** | DENV-2/Asian2 |
| DV-3 F2 | AACCGTGTGTCAACTGGATCAC | DENV-3 |
| DV-3 R2 | TGGCCGTTCARCAATCCT | DENV-3 |
| DV-3 P2 | ***FAM***-TGGCGAAGAGATTC-***MGB*** | DENV-3 |
| DV-4 F3 | GCTTGGAAGCATGCTCAGAGA | DENV-4/gI/IIa/IIb/III |
| DV-4 R3 | GCGCGAATCCTGGGTTT | DENV-4/gI/IIa/IIb/III |
| DV-4 P3 | ***FAM***-TAGAGAGCTGGATACTCA-***MGB*** | DENV-4/gI/IIa/IIb/III |
| DV-4 GS F | CATGGAAACATGCCCAGAGA | DENV-4 GS |
| DV-4 GS R | CCAAGAGTGCAAATCCTGGATT | DENV-4 GS |
| DV-4 GS P | ***FAM***-TGGAGAGCTGGATACTTA-***MGB*** | DENV-4 GS |

Reporter dye (FAM) and quencher (MGB) elements are indicated in bold and italics

#### Phylogenetic analysis

Nucleic acid of dengue complete coding sequences were obtained from Genbank or sequenced in our lab and/or by the European virus archive. Dengue genome were aligned using MEGA software. Best model was determined for each alignment and phylogenetic reconstruction was performed by maximum likelihood using MEGA software with 500 bootstaps.

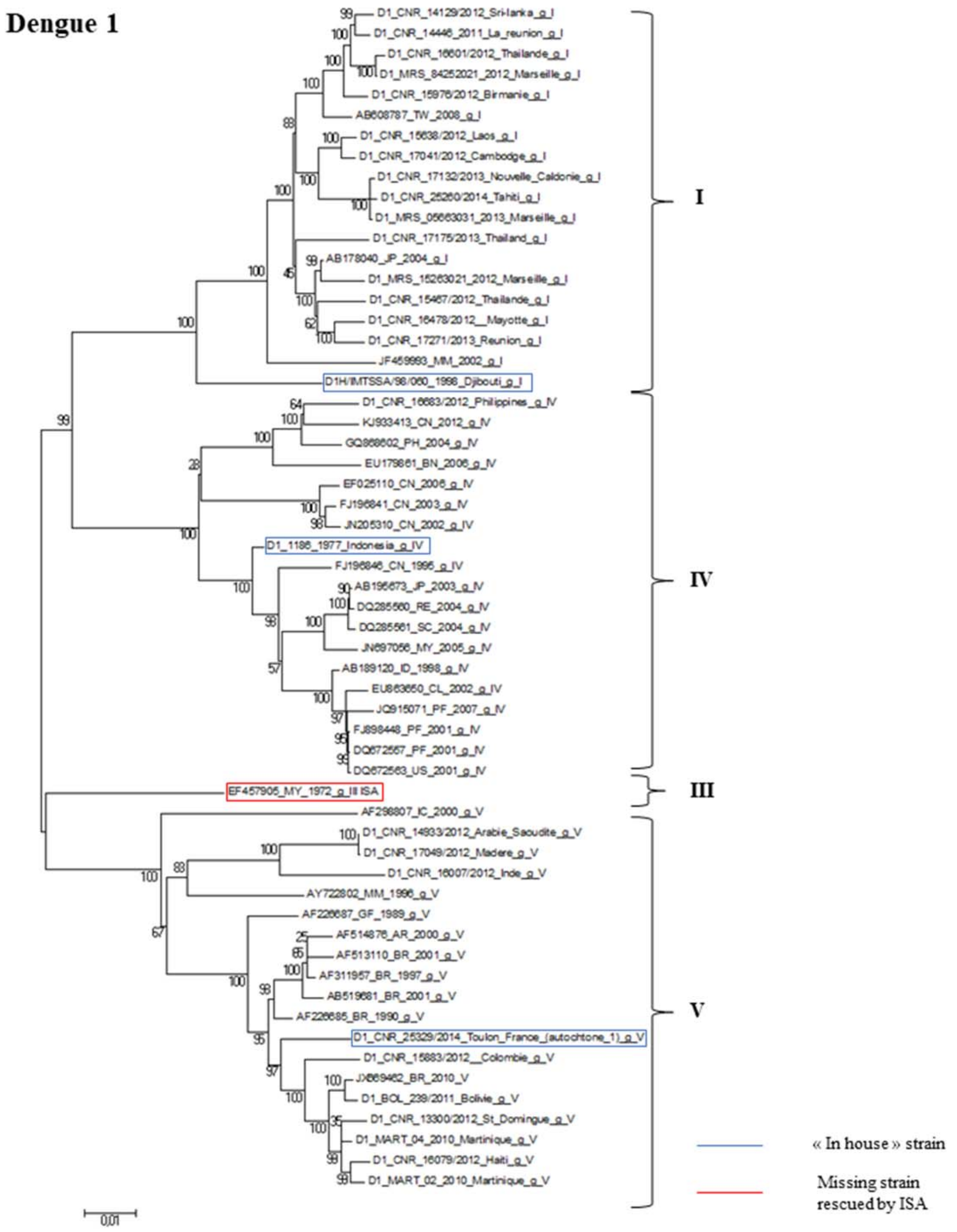


**Supplemental figure 1: reduced maximum likelihood phylogenetic tree (GTR+G model with 500 bootstraps) of the dengue 1 serotype using full nucleic acid coding sequence.** Sequenced selected in the panel are highlighted in blue for the clinical strain, in red for the ISA produced strains.

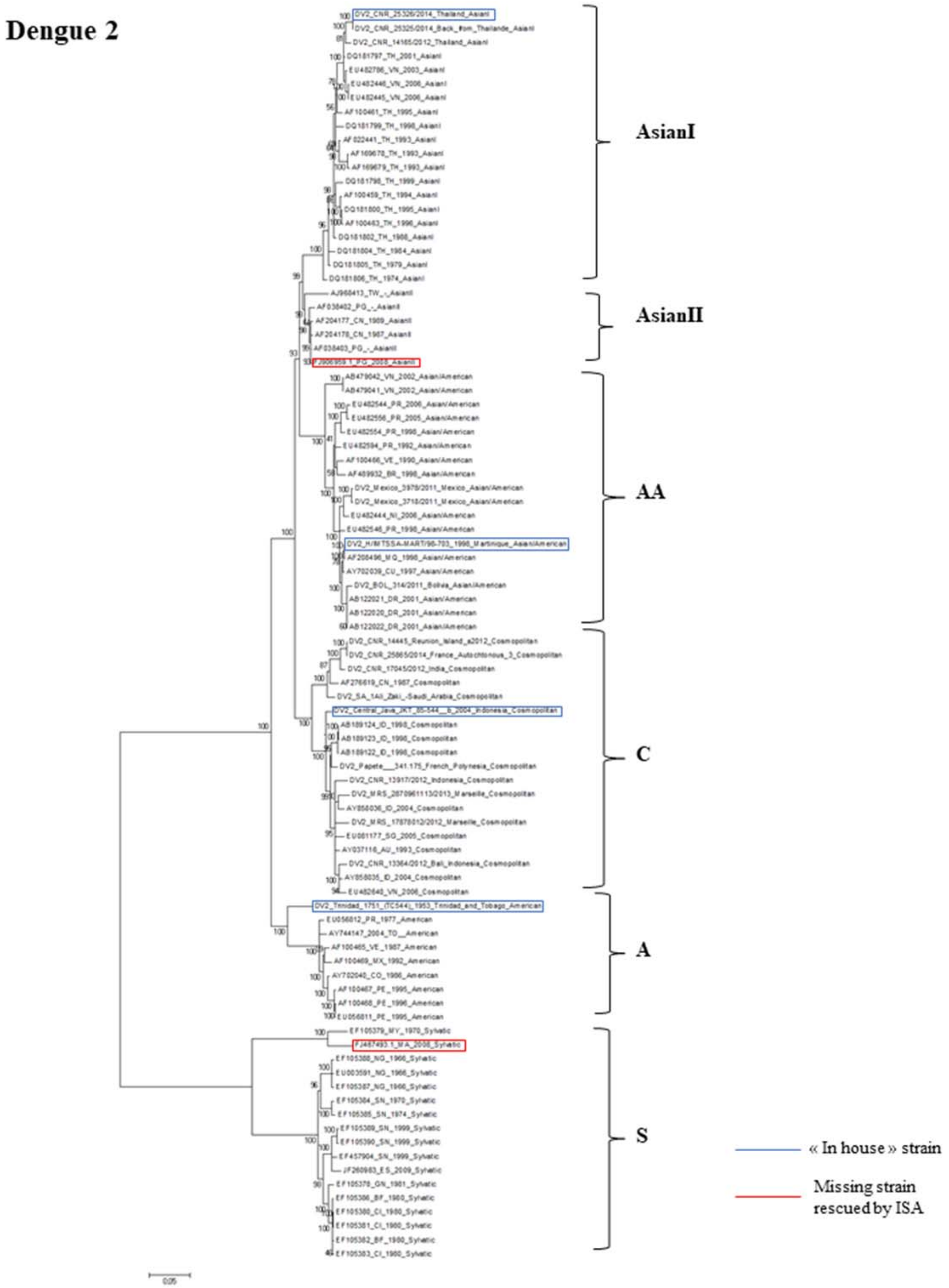


**Supplemental figure 2: reduced maximum likelihood phylogenetic tree (GTR+G+I model with 500 bootstraps) of the dengue 2 serotype using full nucleic acid coding sequence.** Sequenced selected in the panel are highlighted in blue for the clinical strain, in red for the ISA produced strains. AA: Asian American, A: American, C: cosmopolitan, S: Sylvatic.

### Dengue 3

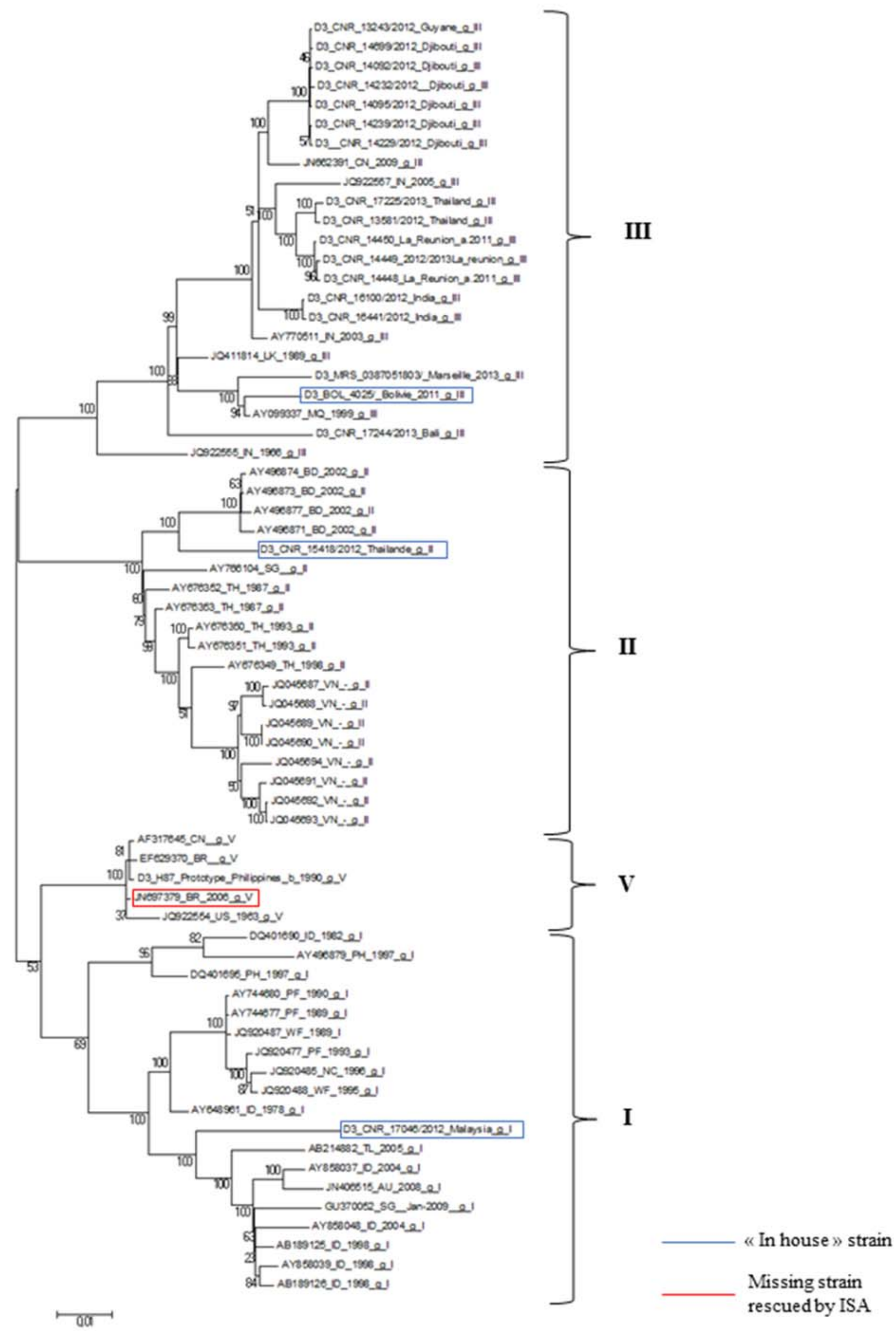


**Supplemental figure 3: reduced maximum likelihood phylogenetic tree (GTR+G+I model with 500 bootstraps) of the dengue 3 serotype using full nucleic acid coding sequence.** Sequenced selected in the panel are highlighted in blue for the clinical strain, in red for the ISA produced strains.

**Dengue 4**

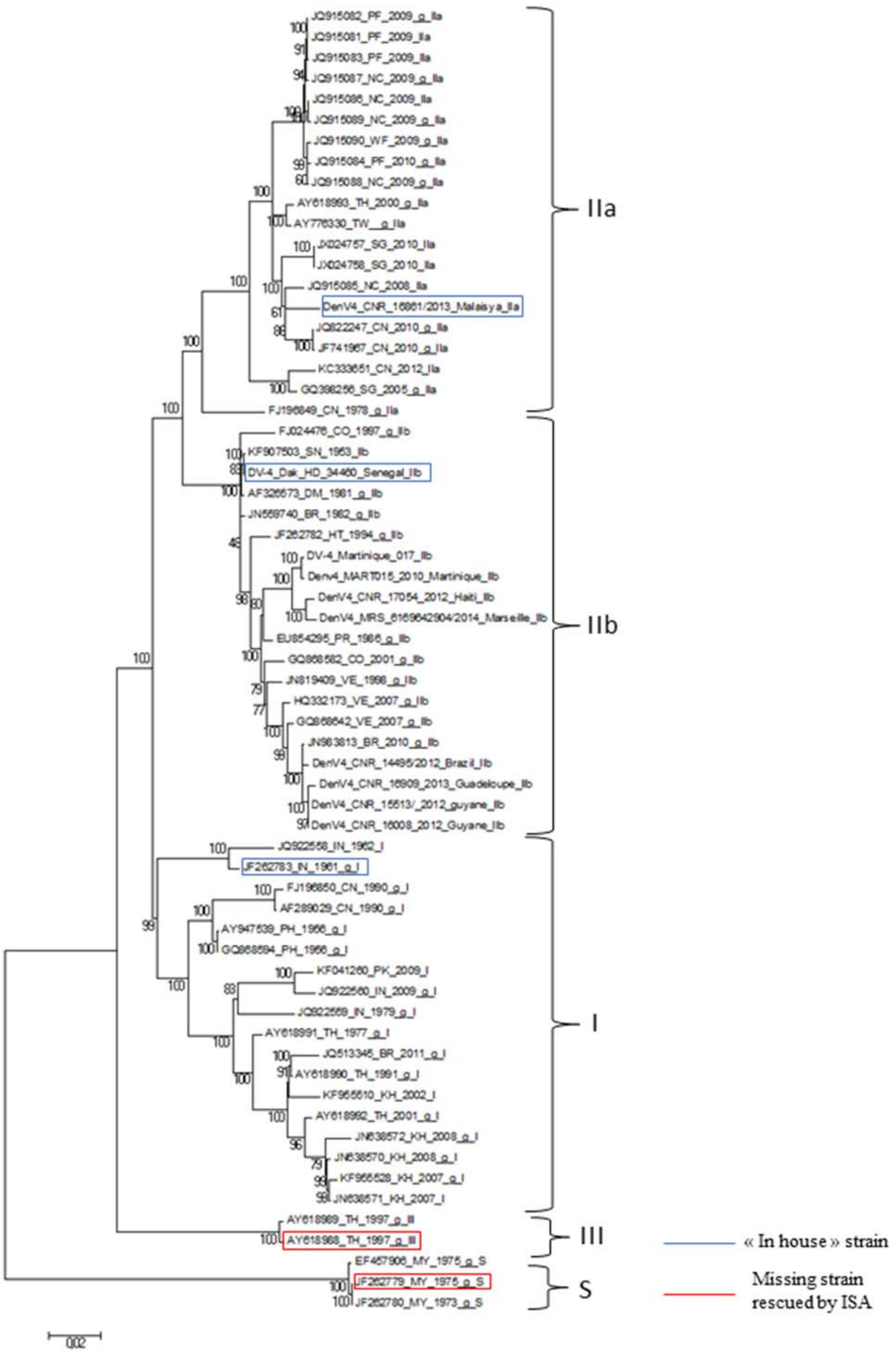


**Supplemental figure 4: reduced maximum likelihood phylogenetic tree (GTR+G+I model with 500 bootstraps) of the dengue 4 serotype using full nucleic acid coding sequence.** Sequenced selected in the panel are highlighted in blue for the clinical strain, in red for the ISA produced strains. S: Sylvatic
